# Localizing spontaneous memory reprocessing during human sleep

**DOI:** 10.1101/2021.11.29.470230

**Authors:** Lea Himmer, Zoé Bürger, Leonie Fresz, Janina Maschke, Lore Wagner, Svenja Brodt, Christoph Braun, Monika Schönauer, Steffen Gais

## Abstract

Reactivation of newly acquired memories during sleep across hippocampal and neocortical systems is proposed to underlie systems memory consolidation. Here, we investigate spontaneous memory reprocessing during sleep by applying machine learning to source space-transformed magnetoencephalographic data in a two-step exploratory and confirmatory study design. We decode memory-related activity from slow oscillations in hippocampus, frontal cortex and precuneus, indicating parallel memory processing during sleep. Moreover, we show complementary roles of hippocampus and neocortex: while gamma activity indicated memory reprocessing in hippocampus, delta and theta frequencies allowed decoding of memory in neocortex. Neocortex and hippocampus were linked through coherent activity and modulation of high-frequency gamma oscillations by theta, a dynamic similar to memory processing during wakefulness. Overall, we noninvasively demonstrate localized, coordinated memory reprocessing in human sleep.

## Introduction

New memories are stabilized and strengthened through synaptic and systems-level modifications of their physical representations. Replay of neural activity patterns, which is supposed to occur during the upstate of the electrophysiological SO in sleep, has been proposed as a central mechanism by which the hippocampus trains slower-learning neocortical memory networks (Klinzing et al., 2019; Zhang et al., 2018). In rodents, memory processing during sleep can be observed as a sequential replay of learning-related firing patterns (Euston et al., 2007), and optogenetic manipulation of reactivation can steer memory consolidation (de Sousa et al., 2019). In humans, memory reactivation during sleep is more difficult to identify. While it is possible to induce reactivation using memory cues during sleep (Rasch et al., 2007; Schreiner et al., 2018), spontaneous memory processing has only been indirectly observed at a coarse spatiotemporal resolution (Schönauer et al., 2017). We aimed to identify both the locations and the mechanisms of memory reprocessing during sleep in whole-night 275-channel MEG recordings. Before sleep, participants performed a picture-location memory task that featured pictures of either faces or houses. We performed whole-brain source reconstruction of the sleep data and identified SOs, which are the hallmark of slow-wave sleep and which have been proposed to coordinate memory reactivation (Klinzing et al., 2019; Staresina et al., 2015). These SOs were then classified with support vector machines to detect and localize characteristic signs of face or house processing, which indicate memory reactivation (see Fig. 1).

**Fig. 1.**
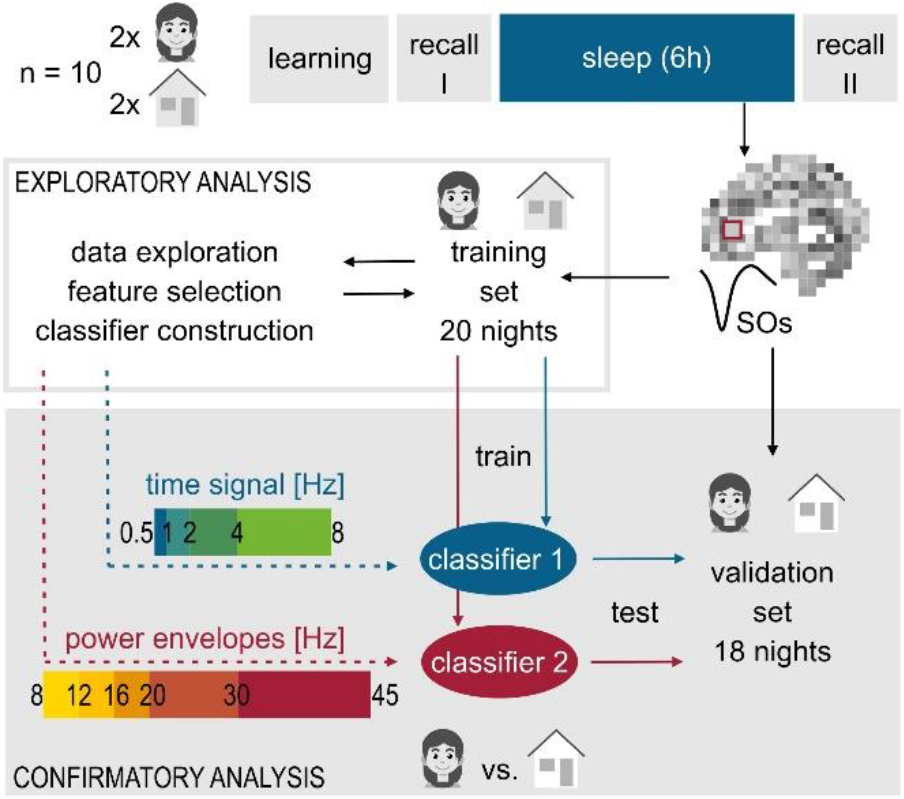
Experimental design and classification procedure. Ten participants stayed in the MEG center for four experimental nights. After memorizing and recalling 100 pictures and their on-screen locations, participants slept in the MEG for 6 hours. In the morning, a second recall took place. Each participant was presented with pictures of houses on two nights and pictures of faces on the remaining two nights. Memory performance did not differ between house and face nights (see Supplementary Fig. 1). Data from 20 nights was used as an exploration set to develop classifiers that could decode information from the sleep slow oscillation. The remaining 18 nights were only used once in a final confirmatory analysis for statistical testing of generalizability. Classification was based on the source space-transformed signal within each voxel of the brain. Two support vector machine classifiers were trained, one was based on the amplitudes of the time-domain signal in slow frequencies (0.5-8Hz), the other was based on the power envelopes of fast frequency oscillations (8-45 Hz), both were time-locked to the downstate of SOs. Statistical significance was determined using cluster-based permutation tests. For the exact classification procedure, see Materials and Methods.

## Results and Discussion

### Memory reprocessing across a distributed fronto-parietal long-term memory network

We found significant memory-related activity in four brain regions: medial prefrontal cortex (mPFC), bilateral inferior frontal gyri (IFG), precuneus and right hippocampus (Fig. 2a, d and Supplementary Table S1). Three of these regions are known as central memory hubs in the brain. The hippocampus is generally recognized as the initial store for episodic memories and is able to acquire new information rapidly, even with single-trial learning (Squire et al., 2004). The role of the mPFC is less clear, although it has been the focus of many previous studies on systems memory consolidation during sleep. It might support encoding and retrieval by providing strategic control processes (Eichenbaum, 2017) or take over indexing functions from the hippocampus after consolidation as proposed by models of systems consolidation (Frankland and Bontempi, 2005). Accordingly, the hippocampus and mPFC are most frequently investigated in the study of systems consolidation and have repeatedly been shown to replay neuronal memory traces during sleep (Euston et al., 2007; Wilson and McNaughton, 1994). It is of particular interest that we can demonstrate neocortical mnemonic activity during sleep in two areas beyond the mPFC – the precuneus and the IFG. While early animal studies provided first hints at a role of the parietal cortex in systems consolidation (Izquierdo and Medina, 1997) and memory replay (Hoffman and McNaughton, 2002) these findings were long overlooked. Only recently, the precuneus, located in the posterior parietal cortex, received attention as the site of a long-term memory engram for declarative memories (Brodt et al., 2018), and it has been shown to interact with the hippocampus in the course of learning and memory consolidation (Hebscher et al., 2019; Himmer et al., 2019). As one of the highest-order association areas, it might store the actual picture-location associations (Gilmore et al., 2015). The fourth region, the IFG, is better known as Broca’s language area, but has also been associated with semantic processing, learning and memory retrieval (Binder et al., 2009). It is active during encoding of subsequently remembered faces (Xue et al., 2010) and during cued reactivation of memories in slow-wave sleep (SWS, Rasch et al., 2007).

**Fig. 2.**
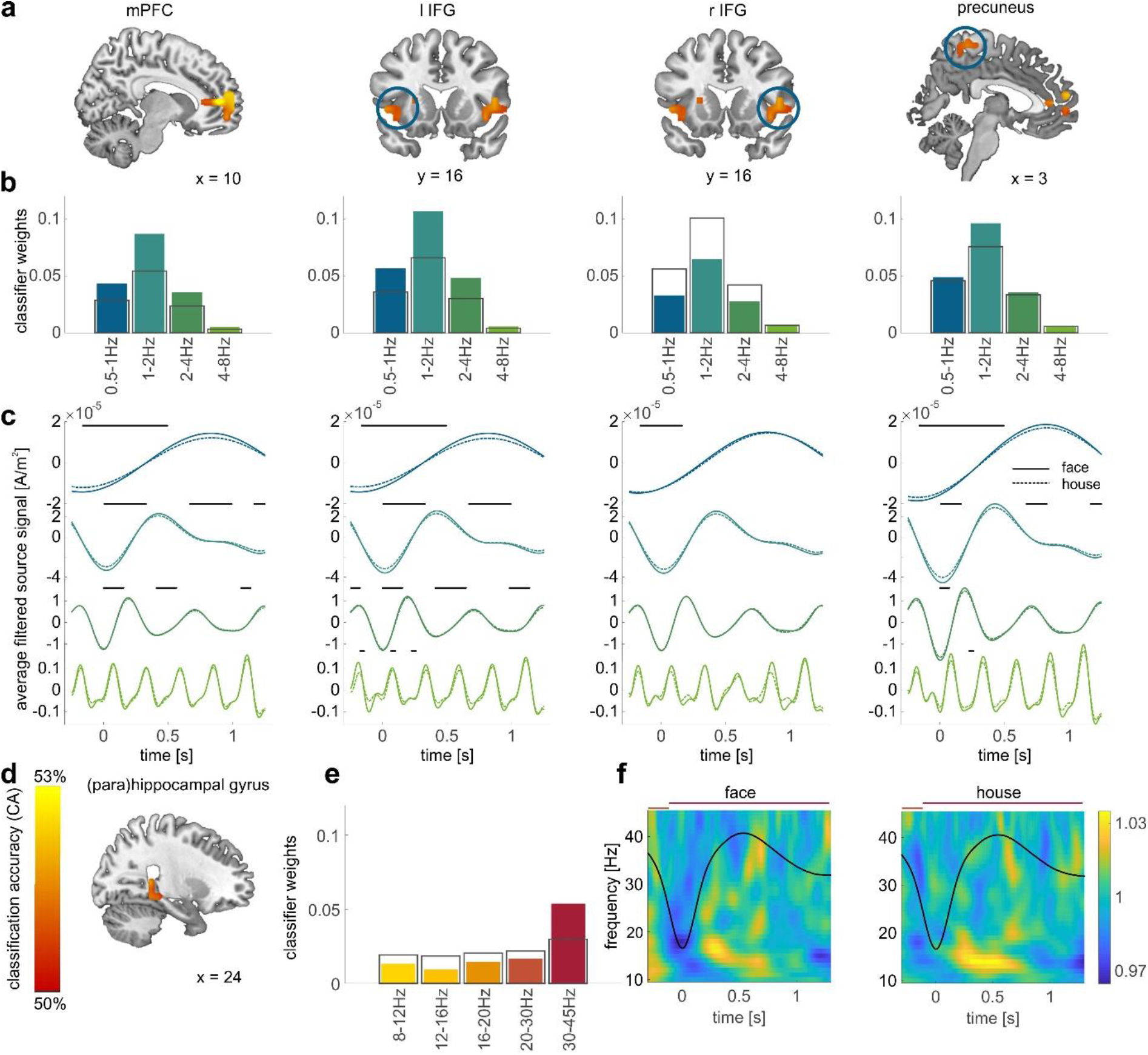
Classification-based information decoding from whole-brain, source space-transformed MEG data. **a)** Analysis of the slow-frequency signal amplitudes, time-locked to the downstate of the slow oscillation. We were able to decode memory-related information from three brain regions: medial prefrontal cortex (mPFC), precuneus and the bilateral inferior frontal gyri (IFG). Brain maps are presented at a cluster-corrected significance level of *p*_corr_≤0.05. Color indicates classification accuracy as shown in d). **b)** We determined the frequency bands that contributed to classification within these clusters by analyzing which classification weights were outside of a 99% confidence interval calculated using permutation statistics. Theta band activity significantly contributed to memory decoding in all four clusters. In mPFC, left IFG and precuneus weights for all slow frequencies below 4Hz were significant (see Supplementary Table S2). Colored bars indicate the average weight of each frequency band, black rectangles indicate the range covered by 99% of permutations, corresponding to *p*_corr_ ≤ 0.04, Bonferroni corrected for multiple comparisons. **c)** A final analysis shows the average time-locked signal filtered in the four frequency bands for face and house conditions. Bars above the curve indicate the time points corresponding to significant classification weights. **d)** Analysis of the fast-frequency power envelopes, time-locked to the downstate of the slow oscillation. Frequencies above 8 Hz allowed classification only in the right posterior hippocampus and parahippocampal gyrus. **e)** Further investigation of classification weights showed that only gamma activity contributed significantly to this classification. Grey rectangles indicate the 99% confidence interval, corresponding to a *p*_corr_≤0.05, Bonferroni corrected for multiple comparisons. **f)** Time-frequency spectra within the hippocampal cluster for face and house nights. The black lines represent the SO in time domain, horizontal bars indicate the intervals during which power classification weights exceed the significance threshold (dark orange 20-30Hz, red 30-45Hz).

Using a multivariate whole-brain classification approach allowed us to look beyond hippocampal-prefrontal interaction and suggests that our models of systems consolidation have to be extended to include a distributed fronto-parietal long-term memory network. Importantly, the use of classifiers enabled us to combine exploratory generation of new hypotheses through repeated training and testing with rigorous statistical confirmation on the independent validation subset without losing statistical power (see Fig. 1 and Methods).

### A role for theta and gamma oscillations in memory reprocessing

Our data goes beyond previous studies by showing that spontaneous memory reprocessing during sleep can involve a whole network of neocortical areas. Additionally, our approach can provide mechanistic insights into how communication between these hubs is achieved. Classification weights reveal that information processing in the hippocampus distinctly differs from that in neocortical areas. Classification of learning material in the neocortex relied on slow activity in the SO (<0.5Hz), delta (0.5-4Hz) and theta (4-8Hz) ranges, slower oscillations, which are regarded as pacemakers of memory reactivation (Klinzing et al., 2019; Staresina et al., 2015). In contrast, classification in the hippocampal area relied on faster gamma (30-45Hz) activity, which is linked to active information transmission (Fig. 2b, e and Supplementary Table S2, Lisman and Jensen, 2013). Looking at the timing of significant classification weights indicates that it is mostly the rising phase of the respective oscillation that carries memory relevant information, particularly in frequencies <4Hz (Fig. 2c, f). Our data is therefore consistent with models of systems consolidation assuming a reactivation of newly encoded information in the hippocampus that provides additional training to neocortical long-term memory networks (Frankland and Bontempi, 2005). This long-range communication must be synchronized to ensure that neurons across the neocortex are in an excitable state when gamma-associated information from hippocampal replay is received. Such synchronization is supposed to rely on rhythmic upstates of the SO, which can coordinate neural firing across long distances (Sirota et al., 2003).

Importantly, our data not only show an involvement of the well-known neocortical SOs in memory reprocessing, but of theta oscillations as well. These have so far been mainly discussed with respect to memory consolidation in REM sleep, during which they dominate brain activity (Klinzing et al., 2019; Nishida et al., 2009). Our findings are in agreement with recent findings on evoked memory reactivation in humans that showed increased theta power following memory cues, which led to gains in memory performance (Schreiner et al., 2015). Within this framework, theta oscillations might enable neocortical plasticity, as they have been linked to synaptic long-term potentiation (Werk and Chapman, 2003). A similar role in memory reactivation has previously been proposed for sleep spindles, which have been consistently found to be temporally coupled to SOs and to support memory consolidation (Gais et al., 2002; Klinzing et al., 2019). Surprisingly, we did not find a significant contribution of spindle frequencies (12-16Hz) with our classification approach. However, we believe that using time-locked amplitudes instead of power envelopes might have yielded a different result, as the precise timing of the spindle within the SO seems to be of importance (Cairney et al., 2018). In sum, our results on classification weights indicate, that the model of a neocortical-hippocampal SO-spindle-ripple cascade of memory replay (Staresina et al., 2015) has to be updated by including a critical role for theta oscillations, which could also be subject to orchestration by the SO.

### Hippocampo-neocortical interactions during memory reprocessing

To further elucidate the differential roles of the four brain regions revealed by classification analysis and their interactions, we investigated two measures of connectivity between neocortical and hippocampal clusters: coherence, which indicates that two brain regions operate at the same frequency in a coordinated fashion, and phase-amplitude coupling, which signifies that the phase of a (slower) oscillation in one area modulates the occurrence and strength of a (faster) oscillation in another region. We found significantly stronger coherence between frontal (right IFG, mPFC) areas and the hippocampus in the low frequency range (<2Hz) during SOs than during random periods before or after (‘non-SO events’; Fig. 3a and Supplementary Table S3). This coherence of SOs could provide the timing for hippocampal memory reactivation and simultaneous integration of information in the neocortex (Sirota et al., 2003; Staresina et al., 2015). When looking at phase-amplitude coupling, we observed the expected delta-spindle coupling between all neocortical areas and the hippocampus, which did not significantly differ between conditions (Supplementary Fig. 3). In contrast, we memory category specific modulation of hippocampal gamma activity by the theta rhythm of the mPFC (Fig. 3b). This finding hints at an organizing role of the frontal cortex and at the hippocampus as the source of information replay, and it indicates that theta oscillations represent a neocortical mechanism for the orchestration of hippocampal high-frequency activity. Consistent with our observations, recent findings on evoked memory reactivation in humans showed theta-phase coordinated memory reactivation, which fluctuated at the SO frequency (Schreiner et al., 2018), which suggests that similar processes underlie both spontaneous and cued reactivation. Theta-gamma coupling has previously been proposed to be an effective neural code of inter-regional network communication and has been associated mainly with memory processing in wakefulness (Köster et al., 2014; Tort et al., 2009). Here, the difference in theta-gamma coupling across memory categories suggests that the neocortex controls which information should be reprocessed in the hippocampus. Finding theta oscillations both to contribute to memory decoding and to influence hippocampal high-frequency activity indicates that its role in memory reactivation during slow-wave sleep deserves further investigation.

**Fig. 3.**
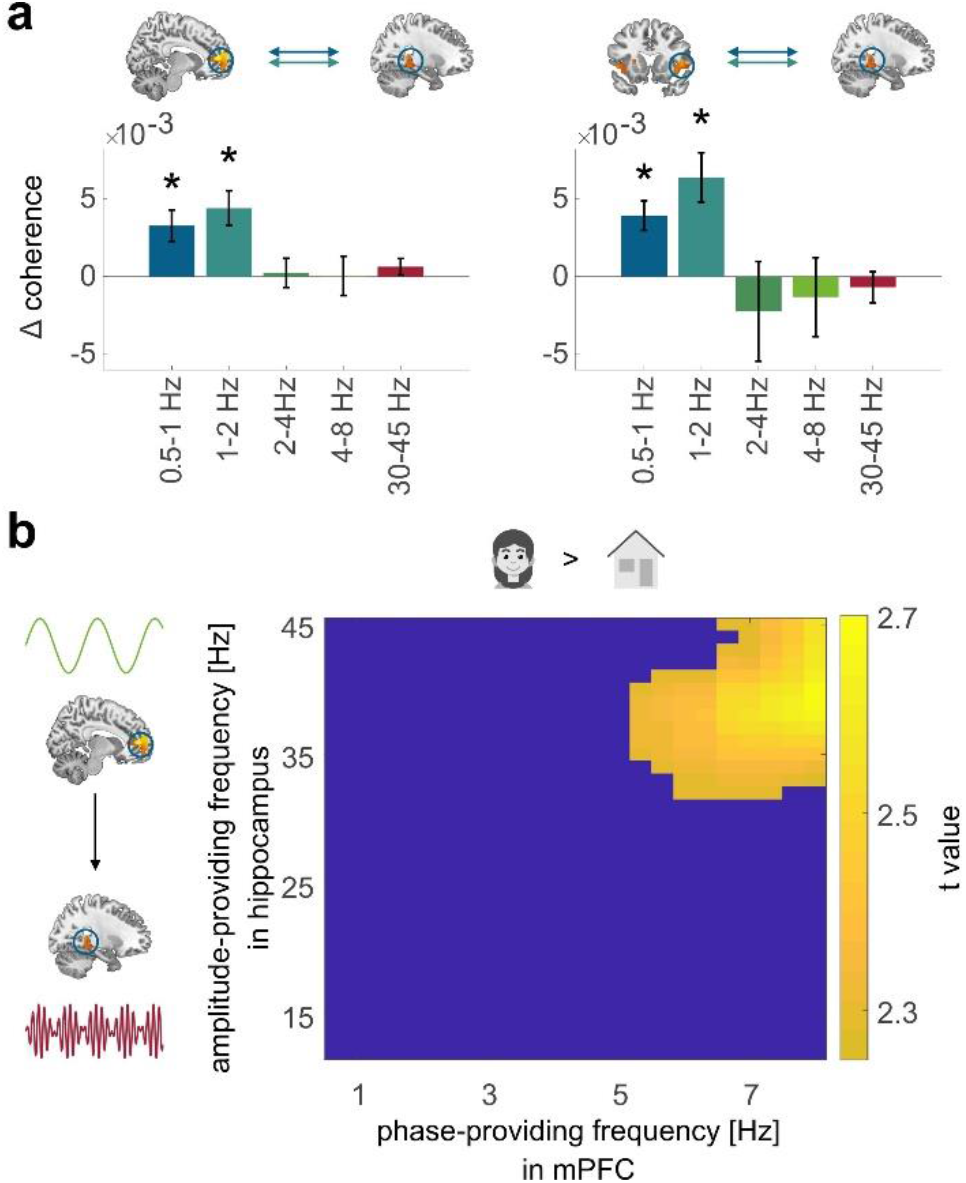
Connectivity between neocortical areas and the hippocampus. a) Coherence between mPFC and hippocampus (left) as well as between right IFG and hippocampus (right) was increased during SOs in the SO and low delta frequencies (< 2Hz) indicating coordinated activity between both areas. Coherence during SOs did not differ from non-SO events for the other frequencies and clusters (see Supplementary Fig. 2 and Supplementary Table S3). b) The phase of mPFC theta oscillations modulated the amplitude of hippocampal low gamma activity. The cluster shown survived cluster correction by permutation testing with p_clust_ = 0.048 (see also Supplementary Fig. 3).

## Conclusions

Overall, our findings demonstrate that using information-based analyses of source space-transformed, tomographic MEG data is a valuable approach to localize cognitive functions in the brain. We present the first evidence of spontaneous memory reprocessing in a network of distinct neocortical regions and the hippocampus during sleep in healthy humans and suggest a role for the posterior parietal cortex alongside the mPFC in memory consolidation. In accordance with recent animal and invasive patient recordings (Helfrich et al., 2019; Todorova and Zugaro, 2019), we demonstrate that memory processing during sleep occurs spontaneously during slow and delta oscillations and is linked to hippocampal gamma activity. Additionally, our data indicate that theta activity, albeit not as apparent during SWS as during REM sleep, also contributes significantly to memory reactivation and plays a central role in joint neocortical-hippocampal memory reprocessing. Analogous to its role in memory during wakefulness, it might support memory reprocessing through the sequential organization and integration of nested hippocampal high-frequency activity, or, it might provide the required theta burst stimulation for LTP induction during SWS, which is otherwise a period of lower potential for synaptic potentiation (Brzosko et al., 2019; Klinzing et al., 2019). The upstate of the sleep slow oscillation therefore seems to use a similar mechanism to organize memory processing as does wakefulness.

## Methods

### EXPERIMENTAL MODEL AND SUBJECT DETAILS

12 young, healthy volunteers took part in the study (8 female; mean age = 25.08 years, sd = 2.64 years). Participants were native German speakers, right handed, and reported to be non-smokers and not to suffer from any sleep or psychological disorders. All participants followed a regular sleep schedule with habitual sleep durations between 6 and 9 hours, as reported in sleep diaries and the Munich Chronotype Questionnaire (Roenneberg et al., 2007). They did not engage in shift work or experience jetlag within 6 weeks of the experiment. Participants were instructed to consume no alcohol and caffeine on experimental days. Experimental procedures were approved by the Ethics Committee of the Medical Faculty of the University of Tübingen. All participants gave informed written consent before participating in the experiment. Data of one subject was excluded because of a missing MRI scan and excessive movement during MEG measurements. Another two subjects aborted the experiment because they were unable to fall asleep in the MEG – one in the adaptation night, one in experimental night 3. Data from experimental night 1 and 2 of this subject were included in analyses. In total, 38 experimental nights from 10 subjects were analyzed in the current study.

## METHOD DETAILS

### General Procedure

Each participant spent 5 nights in the laboratory. The first of these nights served as adaptation night, to get participants accustomed to sleeping in the MEG environment. Participants were asked to go to sleep at their usual bedtime. Sleep was monitored by polysomnography, MEG, a video camera and an interphone connection. Data recorded in these nights was not used further. Subjects were awakened an hour before their usual getting up time.

The first experimental night immediately succeeded the adaptation night, all further experimental nights were spaced 5-8 days apart to allow participants to recover sleep in between sessions. All experimental nights followed the same procedure: Subjects arrived at the laboratory 3 hours before their usual bedtime. Electrodes for polysomnography were attached, before participants performed a learning task for 60-70 minutes in a sitting position in the MEG. Afterwards, the MEG was moved into a horizontal position to allow lying down in supine position. Participants performed a recall task in the MEG and a 10-minute wake resting baseline. Subsequently, the light was turned off and subjects were asked to try and fall asleep. Participants were woken up 6 hours after the first sleep spindles or k-complexes were observed, unless participants were in REM sleep at that time. In this case, participants were woken up after subjects returned to S2 sleep. Participants then left the MEG and were given 30 minutes in order to properly wake up. Afterwards, they returned to the MEG for a second recall and a working memory task. Before leaving, the position of the three fiducial markers and the head shape was a recorded using a 3D positions monitoring systems (Polhemus, Colchester, VT). Participants came back after completing the study in order to acquire a structural MRI of their brain if none was available.

### Learning Task

The learning task consisted of four sets of 100 novel pictures, one for each experimental night. Participants were instructed to remember the image and its position on the screen. In two of the nights, pictures of faces were presented, in the other two nights, pictures of houses were used. The order of face and house nights was randomized across subjects. Faces were taken from the Chicago Face Database (Ma et al., 2015). Houses were taken from German real estate websites. All houses were located outside of Tübingen. All 100 pictures were presented in randomized order in 30 repetitions. Each picture was consistently presented in one of the four corners of the screen, so that its position could be learned. Every picture was preceded by a grey empty frame in the location where the picture would appear and participants were instructed to move their gaze to the center of the frame in order to minimize eye-movements during picture presentation. To make sure, that subjects stayed attentive, this frame was colored blue in 5% of presentations and subjects were instructed to react to the color change by pressing a button. Each repetition of the 100 pictures was followed by a self-paced break, during which subjects were informed of their performance in the attention task during the previous block.

Participants were asked to recall the images once before going to sleep and once again after waking up. During recall all 100 stimuli were presented in random order with 50 additional novel distractor items. For each picture, subjects were asked to indicate whether they recognized the picture, whether they remembered its position and if so, what this position was. Distractor pictures were presented only once during the experiment in order to prevent recognition.

### MEG and polysomnographic recordings

275-channel MEG data was acquired on a CTF275 system (VSM, Vancouver, Canada) at a sampling rate of 1171.9Hz. The head position was continuously tracked by three localization coils marking fiducials on the nasion and the left and right pre-auricular point. We acquired additional polysomnographic data by a synchronized EEG system with 10 electrodes for recording of the EOG (4 electrodes), EMG (2 electrodes placed on jaw muscle), ECG (2 electrodes) and EEG (C3 and C4) to allow sleep scoring and identification of artefacts. The reference was placed on the side of the nose, the ground electrode was placed on the collarbone. Recordings of both MEG and EEG were done in 10-min segments, without gap between recordings. Prior to all analyses, the sensor space signal was 0.1 – 200Hz band-pass filtered and downsampled to 500Hz. This signal was used for classification analyses. Sleep scoring was done by two independent raters according to standard criteria (Rechtschaffen and Kales, 1968) on the basis of polysomnographic recordings (see Supplementary Table S4 for sleep staging). Additionally, one rater identified artifact and eye-movement free episodes of REM sleep, which were used as a baseline for determining thresholds for the detection of slow waves.

### General Analysis Rationale

The general rationale of our analysis approach is characterized by four key components. 1) We achieved an unbiased, whole-brain source reconstruction of whole-night data through sLORETA (Pascual-Marqui, 2002), which enables us to explore the feature space of the MEG signal with high temporal resolution and a reasonable spatial resolution in the centimeter range (Gross, 2019). Compared to surface EEG or MEG, source space-transformed data is no longer affected by long-range volume conduction effects. 2) We used a physiologically meaningful event, the SO, to time-lock our analysis. Thus, we had a sufficiently large number of trials for training our classifier, and we could use the time course of signal amplitudes or power envelopes of individual trials for our analysis. Machine learning classification is ideally suited for multi-dimensional data analysis because it does not rely on testing all potential individual univariate hypotheses, and it takes into account the conjoint effects of feature combinations (Haynes and Rees, 2006). However, machine learning and multivariate pattern analysis can be vulnerable to overfitting the classifier to a finite dataset when searching for informative features and optimizing analysis parameters in an exploratory fashion. 3) Therefore, we employed a combination of an exploratory approach and confirmatory statistical testing. It allowed us to perform exploratory analysis with a 2-fold cross-validation on one half of the dataset, in order to identify meaningful features in the recorded signal. The final classifiers were trained on this initial half of the data and validated on the second half. We could therefore use all available information and the maximal statistical power in the final validation tests. Additionally, this procedure greatly reduces problems of multiple testing, because generalization of classification outcomes is tested only once. 4) To determine the significance of classifier outcomes, we used permutation statistics, which is able to provide accurate results even if the null distribution for classification accuracies is unknown (Jamalabadi et al., 2016). To further characterize our multivariate findings, cluster-based permutation testing was used (Maris and Oostenveld, 2007).

### Source localization

We employed whole brain source reconstruction of the signal in this study. First, we constructed a regular 8mm grid in MNI space covering all voxels within the brain. This grid was warped to the individual anatomy of single subjects to enable the reconstruction of sources corresponding to roughly the same region across subjects. We fit a single-shell realistic head model for each subject based on the segmented structural MRI scan, which was warped to MEG space based on the location of the fiducials and the digitized head shape.

Due to the long scanning period, we needed to take changes in head position into account. While most subjects stayed very still, some subjects showed sleep-typical patterns of movement, consisting of long constant head positions with fast shifts in between. In order to distinguish the stable periods between shifts, we implemented a clustering algorithm, which worked as follows: We downsampled headcoil signals to 1Hz and calculated the circumcenter of the three coils to infer head shifts. Head rotation was not used for clustering, as it never exceeded 2.5°. The shifts were entered into an agglomerative hierarchical cluster model, which distinguished head positions that were separated by a distance criterion of 10mm or more. We then calculated the mean head position for these clusters. Only positions, in which subjects stayed for at least 5 min were considered for further analysis. Time intervals that did not correspond to any stable position or corresponded to movements were excluded from further analysis.

Leadfields and spatial filters were calculated separately for each of the stable head positions. We used sLORETA as implemented in the FieldTrip toolbox (Oostenveld et al., 2011) with a regularization parameter lambda of 1.0×10-15 (based on the optimization procedure for lambda used by NUTMEG (Dalal et al., 2011)) to construct filters for each voxel (8mm isotropic). We then projected the signal along the dipole direction that explained the most variance by determining the largest temporal eigenvector of the bandpass filtered signal (0.1-40Hz) through singular value decomposition.

### Artefact detection

Time intervals containing artefacts were detected for all nights. Jumps were detected in all channels on the sensor level with FieldTrip (ft_artefact_jump) with a median filter. The intervals containing artefacts in any sensor were marked as artefacts for all voxels in source space. Muscle artefacts were detected in the EMG channels using ft_artefact_muscle on the Hilbert transform of the band-passed high frequency signal (110-140Hz) and marked as artefacts for all voxels. To account for movements, we excluded all intervals that did not belong to one of the stable head position clusters (see above).

### Slow wave detection

Slow waves were detected in each voxel separately. First, the sensor signal was filtered in the frequency range between 0.3 and 3Hz before being multiplied with the spatial filters corresponding to the voxel position. We then marked local minima and maxima in the signal. The difference between a minimum and the succeeding maximum was defined as the amplitude of a slow wave. While the habitual threshold for detecting slow waves in the EEG is usually 75μV, no such criterion exists for MEG recordings. We therefore used an amplitude criterion based on (eye movement-free) REM sleep, where no slow waves occur. This was done as follows: First, we band-pass filtered the signal from all REM sleep episodes that were visually marked as being free of eye movements and other artefacts in the range between 0.3 and 3Hz, marked local minimal and maxima, and calculated their amplitudes as for the slow waves above. In an iterative process, we then examined the 95 to 99.9 percentile of these amplitudes in steps of 0.1%. In order to exclude an influence of residual artefacts, which would lead to excessively high thresholds, we stopped the procedure when the step to the next higher percentile led to an amplitude increase by more than 20% and used the prior percentile as amplitude threshold. All slow waves that exceeded this criterion and did not overlap with an artefact were then marked as slow waves. The local minimum indicated the downstate and the local maximum indicated the upstate. SO events, which were used for classification analyses, used 2-s intervals of the source space-transformed signal of each voxel starting 0.5 seconds before the downstate peak. We explicitly aimed at investigating slow oscillations (SOs) during deep slow-wave sleep and excluded smaller slow waves and K-complexes. We therefore only considered slow waves that occurred in trains of at least 5 consecutive slow waves as SOs. Detected SOs were most frequent in frontal regions and the lowest number of events occurred in the occipital cortex (see Supplementary Fig. 4).

## QUANTIFICATION AND STATISTICAL ANALYSIS

### Classification and statistical testing

The development of a classifier requires exploration of data in order to identify suitable features. Multiple rounds of feature exploration and classification on the same data set, however, leads to overfitting and an increased false positive rate. Therefore, we explored the data using repeated cross-validation on a training/test set consisting of 20 nights from 10 subjects (1 face and 1 house night each). After we were sufficiently confident that the classifier was optimally adapted to our task, we performed a final confirmatory analysis. The classifier was trained on the 20 nights of the training set and generalization was confirmed with a validation set consisting of 18 nights from 9 subjects (1 face and 1 house night each, see Figure 1), which had not been used during exploration.

The exploratory phase of analysis indicated that information from slower frequencies up to the theta range and faster frequencies can be best used separately by two classifiers. Classifier 1 used the time-domain signal of the lower frequencies. The MEG signal was band-pass filtered using FFT-filters into four separate bands (SO: 0.5-1Hz, low delta: 1-2Hz, high delta: 2-4Hz, theta: 4-8Hz). Because support vector machines (SVMs), which we used for classification, benefit from a sparse feature space, we downsampled the filtered signal to 1.5 times the Nyquist frequency of the upper limit of each band (i.e. 6, 12, 24 and 48 samples, respectively, per band for the 2-s SO interval), leading to a classifier with a total of 90 features. Classifier 2 used the power of frequencies above 8Hz. The signal was band-pass filtered in five separate bands (alpha: 8-12Hz, sigma: 12-16Hz, low beta: 16-20Hz, beta: 20-30Hz, gamma: 30-45Hz). The envelope of each band was sampled at 5Hz, resulting in a classifier with 50 features.

For the final confirmatory analysis, both classifiers were applied according to following pipeline. Classification was done for each voxel of the brain separately, and it included as trials the SOs detected in that voxel signal. To achieve a stable estimate of classification accuracy (CA), classifier training was repeated on all possible subsets of 6 subjects (12 nights) out of 10 subjects, with 24 randomly drawn SO events per night. Training data was z-standardized per feature and used to train an SVM (fitclinear, Matlab 2018b, The Mathworks Inc., Natick, Mass). Validation data was standardized with the means and standard deviations derived from the training set. Classification accuracies were calculated for each training repetition by averaging single-subject accuracies obtained by dividing the number of correctly predicted trials by the total number of trials for this subject. Overall CA was obtained by averaging over all subset repetitions. Participants were only included in the classification when at least 35 SO events were detected in this voxel in both the face and the house night.

Statistical significance was determined by permutation tests in each voxel followed by a non-parametric cluster-based correction for multiple testing. In 500 permutations per voxel, the classification procedure was performed exactly as described above after class labels were randomly permuted across nights. P-values were calculated by comparing the true CAs with the null distributions obtained by permutation. Importantly, permutation was not done on the level of individual events, but trials within nights and nights of participants were kept together in order to preserve the influence of these subclasses on CAs (Jamalabadi et al., 2018; Winkler et al., 2015). Thus, all events from the two nights of each subject in the testing set either kept the correct labels or all labels were flipped. The same was done for permutation of the validation set.

Because we performed classification in all 3203 voxels of the brain, we employed a non-parametric cluster-based Monte Carlo correction to account for multiple testing (based on (Maris and Oostenveld, 2007)). CAs cannot be interpreted without reference to the underlying null-distribution (Jamalabadi et al., 2016). In particular, CAs from different classifiers (e.g. different voxels) cannot be compared directly. We therefore used statistical p-value maps based on voxel-wise permutation tests to compare voxels. To determine significant clusters, we first detected clusters of neighboring voxels with p-values of p≤.05 (using spm_bwlabel, SPM, FIL, London). The product of all p-values within a cluster was saved as the cluster-specific statistic. To obtain the null distribution, we repeated this procedure 1000 times using p-values derived by randomly drawing from the permutation distributions for each voxel. The minimal cluster-specific statistic value was saved for each repetition. The actual cluster statistics were then compared against this null distribution. All clusters based on the real data with a cluster-specific statistic smaller than or equal to the 5th percentile of this null distribution were considered significant, which corresponds to a whole-brain corrected p-value of p_clust_≤0.05.

In addition to classification accuracy, we analyzed feature weights of our linear SVM models, which can give insights into whether a feature carries information. While a feature weight close to 0 indicates little to no information, a high negative or positive weight indicates, that a feature contributes to the distinction between conditions. Feature weights were saved for the classification as well as for all permutations for each voxel. This enabled us to compare feature weights to a null distribution derived by permutation and compute significances.

Feature weights were averaged over all repeated classifications and over all voxels within each significant cluster. We then summarized all weights within a frequency band by averaging the absolute value of all feature weights belonging to the frequency. This results in one weight per frequency band and cluster. We then obtained a p-value by comparing the true weight value with the null distribution of weights obtained from permutation testing. This p-value was multiplied by the number of frequency bands (4 for classifier 1, 5 for classifier 2) to account for multiple comparisons. A value of p_corr_≤0.05 was considered significant.

### Connectivity measures

We calculated the coherence and the modulation index (Tort et al., 2009), between all 4 neocortical clusters given by classifier 1 and the (para)hippocampal cluster given by classifier 2. Coherence was assessed between each neocortical cluster voxel with each (para)hippocampal voxel. We derived Fourier spectra of all SO events within a neocortical voxel for frequencies between 0.5 and 45Hz with a bin width of 0.5Hz. Coherence was then assessed for all frequency bands that showed significant contributions to classification (SO, slow delta, high delta, theta, low gamma). Coherence values were averaged over all hippocampal cluster voxels and all voxels within each neocortical cluster. They were further averaged over all frequencies within each frequency band, resulting in one value per frequency band, night, and neocortical cluster. We performed the same analysis for non-SO events, which were defined by a random offset between 7 and 30 seconds from true SOs. Non-SO events were taken from non-REM sleep and did not include artifacts. Coherence was then compared between SO events and non-SO events by paired t-tests. P-values were multiplied by 5 to account for multiple testing in five frequency bands and considered significant at p_corr_≤0.05.

Additionally, we calculated the modulation index, which is a form of phase-amplitude coupling, between slow frequencies (0.6-8Hz) in neocortical clusters and fast frequencies in the (para)hippocampal cluster (12-45Hz). In a first step, we calculated time-frequency spectra based on SO events in each neocortical voxel using Morlet wavelets for frequencies from 0.6 to 8Hz in steps of 0.33Hz with wavelet widths between 0.2 and 2.5. The same events were used to calculate time-frequency spectra in hippocampal cluster voxels for frequencies between 12 and 45Hz in steps of 1Hz with wavelet widths of 6. Based on these spectra we calculated the modulation index between all neocortical cluster voxels and all (para)hippocampal cluster voxels. Modulation index values were then averaged over all (para)hippocampal voxels and all voxels within each neocortical cluster.

In order to compare modulation index between the face and house condition, we averaged modulation indices over the 2 nights of each condition within each subject. This results in one modulation index matrix per condition, subject, and neocortical cluster. We compared conditions within each neocortical cluster by paired t-tests. To determine cluster-based corrected significances, we then applied an initial threshold of p≤0.05 and obtained clusters within our cross-frequency space. We summed up t-values within each cross-frequency cluster and compared them to the highest summed t-values obtained by 10000 permutations of the data to obtain p-values (Maris and Oostenveld, 2007). This p-value was then multiplied by 2 to account for two-sided testing and considered significant at p_clust_≤0.05.

## Acknowledgments

We thank Jürgen Dax for technical assistance at the MEG centre.

## Author contributions

LH, SB, CB, MS and SG designed the experiments, LH, ZB, JM, and LW acquired the data, and LH, ZB, LF, JM, and LW analyzed the data, LH and SG wrote the manuscript, all authors revised the manuscript.

## Notes

### Competing Interest Statement

The authors have declared no competing interest.

