## Supplementary Information for "Localizing spontaneous memory reprocessing during human sleep"

**Contents**

- Supplementary Figures S1 to S4
- Supplementary Tables S1 to S4

### Supplementary Figures

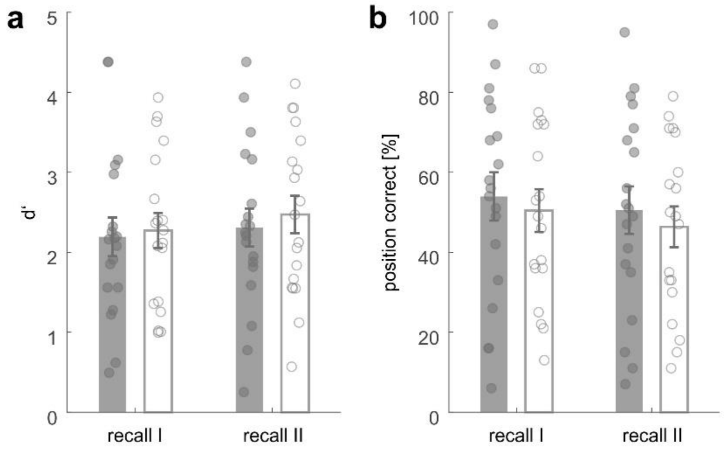


**Fig. S1 Memory performance**. a) The sensitivity index d', a measure of recognition performance, and b) the percentage of pictures for which the correct location was recalled are given for face nights (filled bars) and house nights (open bars) in recall I and recall II. Error bars represent the standard error of the mean, each circle represents data for a single night. Memory performance was assessed by calculating d' and the percentage of pictures that were assigned the correct position after being correctly identified as "old". Values were averaged across nights belonging to the same memory category within subjects. We then performed a repeated measures ANOVA with within-subject factors recall I/recall II and face/house on both measures. For d', we observed an increase in performance from recall I to recall II (main effect $F_{1,9} = 7.601$, $p = 0.022$). This indicates, that participants were better in distinguishing old and new pictures in the morning after sleep compared to the evening. In contrast, we observed a concurrent decrease in position recall overnight (main effect $F_{1,9} = 35.053$, $p \leq 0.001$). Both d' and position recall showed no difference between memory categories (d': main effect face/house $F_{1,9} = 0.359$, $p = 0.564$, interaction $F_{1,9} = 0.909$, $p = 0.365$; position recall: main effect face/house $F_{1,9} = 0.681$, $p = 0.431$, interaction $F_{1,9} = 0.364$, $p = 0.561$).

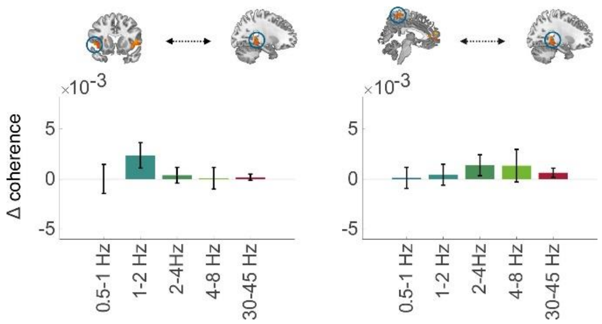


**Fig S2. Coherence between left IFC, precuneus and hippocampus.** Coherence between left IFG and hippocampus (left panel) did not show significant changes during SO events compared to non-SO. Similarly, we did not observe changes in coherence during SOs between precuneus and hippocampus (see Supplementary Table S3 for statistics).

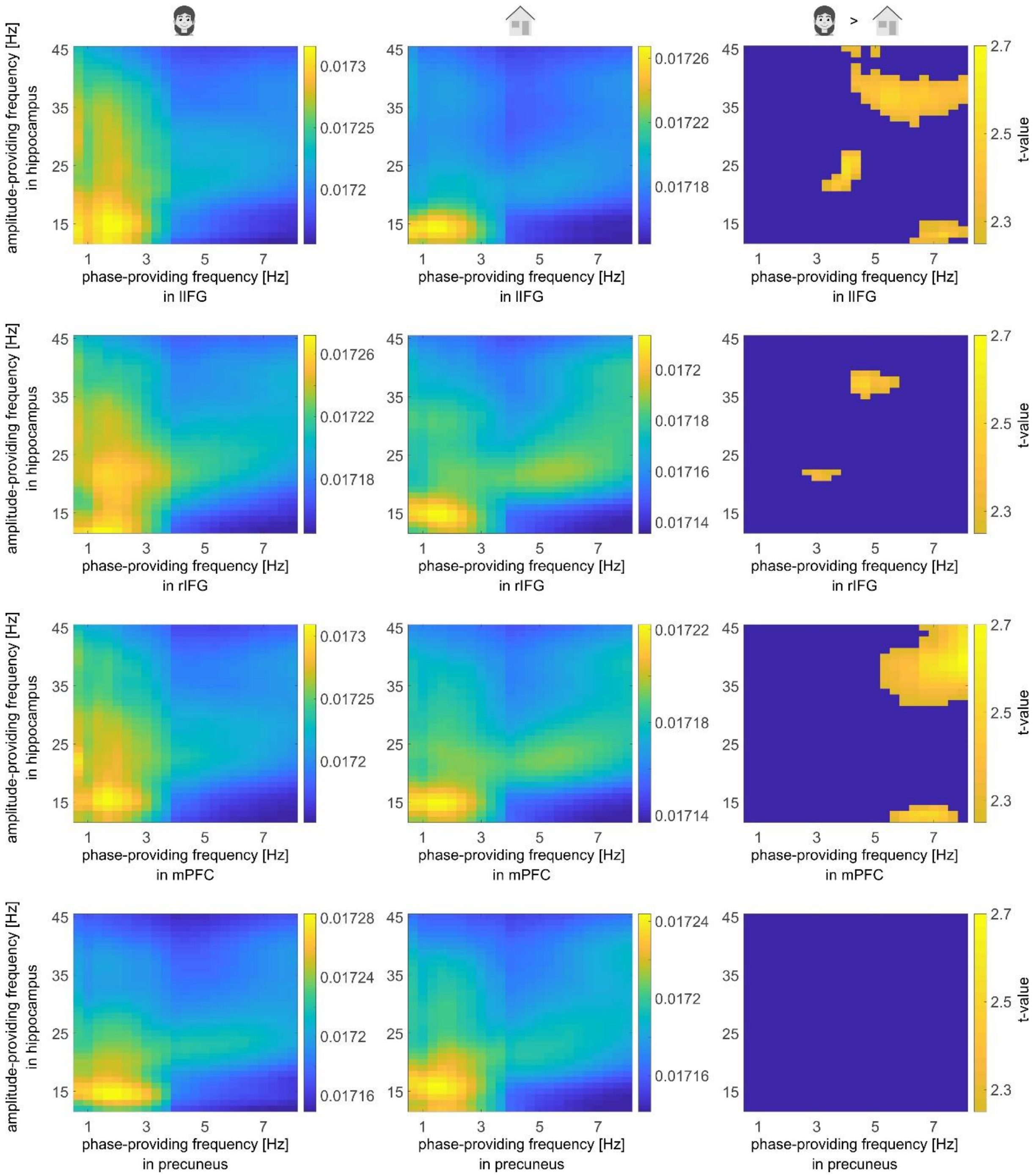


**Fig S3 Modulation index between all neocortical clusters and right (para)hippocampus.** Shown are the average modulation matrices across subjects and conditions (left column: faces, middle column: houses). The right column depicts the result of a paired t-test between conditions at an uncorrected level of $p \leq 0.05$. Please note, that only the theta-gamma coupling cluster between mPFC and hippocampus survived cluster-correction by permutation tests.

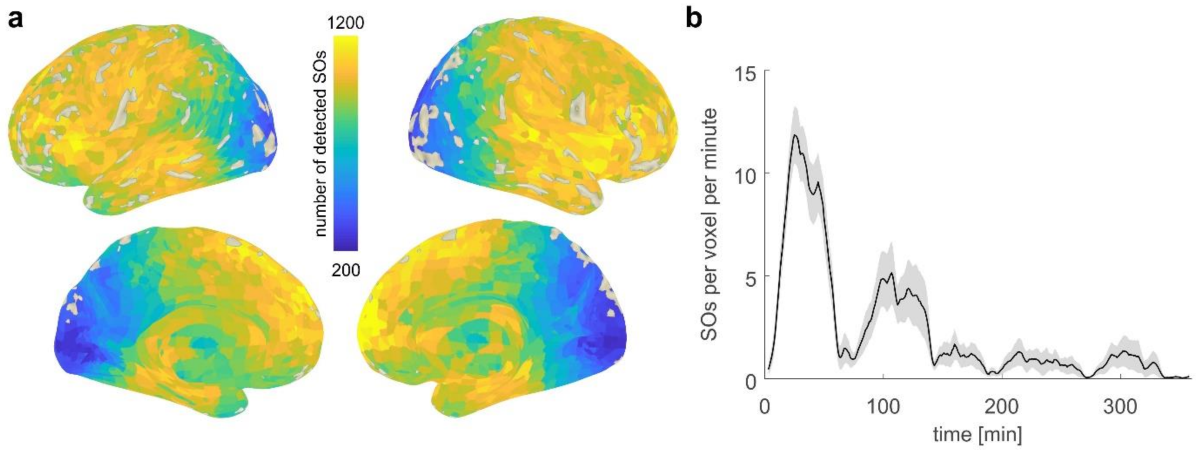


**Fig. S4 SO detection** a) Topography of detected SOs. The average number of SOs over all 38 nights per voxel is shown in a surface projection (top: lateral view, bottom: medial view). Most SOs were detected in frontal regions, while the lowest number of SOs was detected in occipital cortex. b) Number of SOs detected per minute per voxel over all nights. Time 0 corresponds to the first slow wave detected within a night. The number of SOs per minute and voxel was assessed per night by counting all SOs within 5-min windows that were shifted by 1 minute, and dividing by the number of voxels within the brain. The line is the average time course; the shaded area indicates the standard error of the mean.

### Supplementary Tables

#### Table S1.

**List of brain regions that allowed classification.** The table denotes all clusters that survived cluster correction by permutation testing at $p_{clust}$≤0.05. Coordinates are given as the median of all voxel coordinates within a cluster. # indicates the number of voxels within a cluster.

| anatomical regions | | MNI coordinates (mm) x | y | z | # | max CA | $p_{clust}$ |
|---|---|---|---|---|---|---|---|
| **Classifier 1** | | | | | | | |
| left | Inferior frontal gyrus | -40 | 32 | 0 | 19 | 0.527 | ≤0.001 |
| | Insula | | | | | | |
| right | Inferior frontal gyrus | 48 | 16 | 0 | 6 | 0.522 | 0.033 |
| | Insula | | | | | | |
| right | Superior frontal gyrus | 8 | 56 | 8 | 24 | 0.545 | ≤0.001 |
| | Anterior cingulum | | | | | | |
| | Medial frontal gyrus | | | | | | |
| left | Anterior cingulum | | | | | | |
| | Superior frontal gyrus | | | | | | |
| | Medial frontal gyrus | | | | | | |
| left | Precuneus | -4 | -40 | 64 | 10 | 0.525 | ≤0.001 |
| | Paracentral lobule | | | | | | |
| **Classifier 2** | | | | | | | |
| right | Hippocampus | 24 | -40 | -4 | 8 | 0.521 | 0.004 |
| | Parahippocampal gyrus | | | | | | |

**Table S2.**

**Statistics on classifier weights**. The table denotes p-values derived by comparing classification weights with randomization weights. P-values were corrected for the number of comparisons done per region and are considered significant at $p_{corr} \leq 0.05$, indicated by an asterisk.

| anatomical regions | | $p_{corr}$ | | | | |
|---|---|---|---|---|---|---|
| **Classifier 1** | | **0.5-1 Hz** | **1-2 Hz** | **2-4Hz** | **4-8Hz** | |
| left | Inferior frontal gyrus | ≤0.008* | ≤0.008* | ≤0.008* | ≤0.008* | |
| right | Inferior frontal gyrus | 0.583 | 0.439 | 0.408 | 0.016* | |
| | Medial prefrontal cortex | ≤0.008* | ≤0.008* | ≤0.008* | ≤0.008* | |
| left | Precuneus | 0.024* | 0.016* | 0.032* | 0.032* | |
| **Classifier 2** | | **8-12Hz** | **12-16Hz** | **16-20Hz** | **20-30Hz** | **30-45Hz** |
| right | Hippocampus | ≥1 | ≥1 | ≥1 | 0.449 | ≤0.01* |

**Table S3.**

**Statistics on coherence analyses of neocortical clusters to the right hippocampal cluster.** The table denotes statistics obtained by comparing coherence in the specified frequency bands during SOs with coherence during non-SO events. P-values were corrected for 5 comparisons per region and are considered significant at $p_{corr} \leq 0.05$, indicated by an asterisk.

| | | Frequency band | | | | | | | | | |
|---|---|---|---|---|---|---|---|---|---|---|---|
| | | 0.5-1Hz | | 1-2Hz | | 2-4Hz | | 4-8Hz | | 30-45Hz | |
| anatomical region | | $t_{37}$ | $p_{Corr}$ | $t_{37}$ | $p_{Corr}$ | $t_{37}$ | $p_{Corr}$ | $t_{37}$ | $p_{Corr}$ | $t_{37}$ | $p_{Corr}$ |
| left | Inferior frontal gyrus | .010 | ≥1 | 1.865 | .351 | .499 | ≥1 | .084 | ≥1 | .613 | ≥1 |
| right | Inferior frontal gyrus | 4.155 | .001* | 3.990 | .002* | -.696 | ≥1 | -.518 | ≥1 | -.687 | ≥1 |
| | Medial prefrontal cortex | 3,234 | .013* | 3.933 | .002* | .253 | ≥1 | .035 | ≥1 | 1.207 | ≥1 |
| left | Precuneus | .119 | ≥1 | .432 | ≥1 | 1.330 | .959 | .826 | ≥1 | 1.442 | .789 |

**Table S4.**

**Sleep stage distribution and total sleep time.** TST: Total sleep time; S1-S2: sleep stage 1 and 2; SWS: slow wave sleep; REM: rapid eye movement sleep; Sleep latency refers to the time from the start of recordings until the first episode of S2 was scored. All values are given as mean ± standard deviation.

| | Time [min] | % of TST |
|---|---|---|
| Time in MEG | 359.65 ± 14.06 | |
| Sleep onset latency | 3.70 ± 2.87 | |
| TST | 350.29 ± 20.45 | |
| Wake | 11.44 ± 19.19 | 3.22 ± 5.27 |
| S1 | 24.05 ± 24.76 | 6.81 ± 6.86 |
| S2 | 163.24 ± 28.47 | 46.61 ± 7.50 |
| SWS | 89.06 ± 23.55 | 25.50 ± 6.79 |
| REM | 61.05 ± 19.56 | 17.45 ± 5.59 |
| movement | 1.44 ± 2.64 | 0.41 ± 0.73 |
